# Metadata retrieval from sequence databases with *ffq*

**DOI:** 10.1101/2022.05.18.492548

**Authors:** Ángel Gálvez-Merchán, Kyung Hoi (Joseph) Min, Lior Pachter, A. Sina Booeshaghi

## Abstract

We present a command-line tool, called *ffq*, for querying user-generated data and metadata from sequence databases. The code can be found here: https://github.com/pachterlab/ffq.

## Introduction

The extraordinary large volume of user-generated sequencing data available in public databases is increasingly being utilized in research projects alongside novel experiments (Simon *et al*., 2018; Razmara *et al*., 2019; Lung *et al*., 2020; Rajesh *et al*., 2021; Hippen and Greene, 2021; Wartmann *et al*., 2021; Kasmanas *et al*., 2021; Huang *et al*., 2021; Klie *et al*., 2021; Booeshaghi *et al*., 2022). Collation of metadata is crucial for such reuse of publicly available data since it can provide information about the samples assayed and can facilitate the acquisition of raw data. For example, *sra-tools* enables users to query and download data from the National Center for Biotechnology Information Sequence Read Archive (NCBI SRA), which currently hosts 13.67 PB of data. An alternative to *sra-tools* is the *pysradb* tool (Choudhary, 2019). *pysradb* was developed to access metadata from the Sequence Read Archive (SRA), using metadata obtained from the regularly updated SRAdb SQLite database (Zhu *et al*., 2013). MetaSRA adds additional standardized metadata on top of the SRAdb SQLite database (Bernstein *et al*., 2017) and also provides an API for accessing them. While these and other tools (Mahi *et al*., 2019; Li *et al*., 2018; Eaton, 2020; Bernstein *et al*., 2020) have proven to be very useful, they are limited in terms of the scope of databases they provide access to. We developed *ffq* to facilitate metadata retrieval from a diverse set of databases, including

1. National Center for Biotechnology Information Sequence Read Archive (SRA) and Gene Expression Omnibus (GEO),
2. European Molecular Biology Lab-European Bioinformatics Institute European Nucleotide Archive (EMBL-EBI ENA),
3. DNA Data Bank of Japan Gene Expression Archive (DDBJ GEA), and
4. Encyclopedia of DNA Elements (ENCODE) database (Davis *et al*., 2018; ENCODE Project Consortium, 2012).

In order to facilitate a modular architecture for *ffq*, we first studied the structure of these databases in detail to identify commonalities and relationships between them (Figure 1).

**Figure 1:**
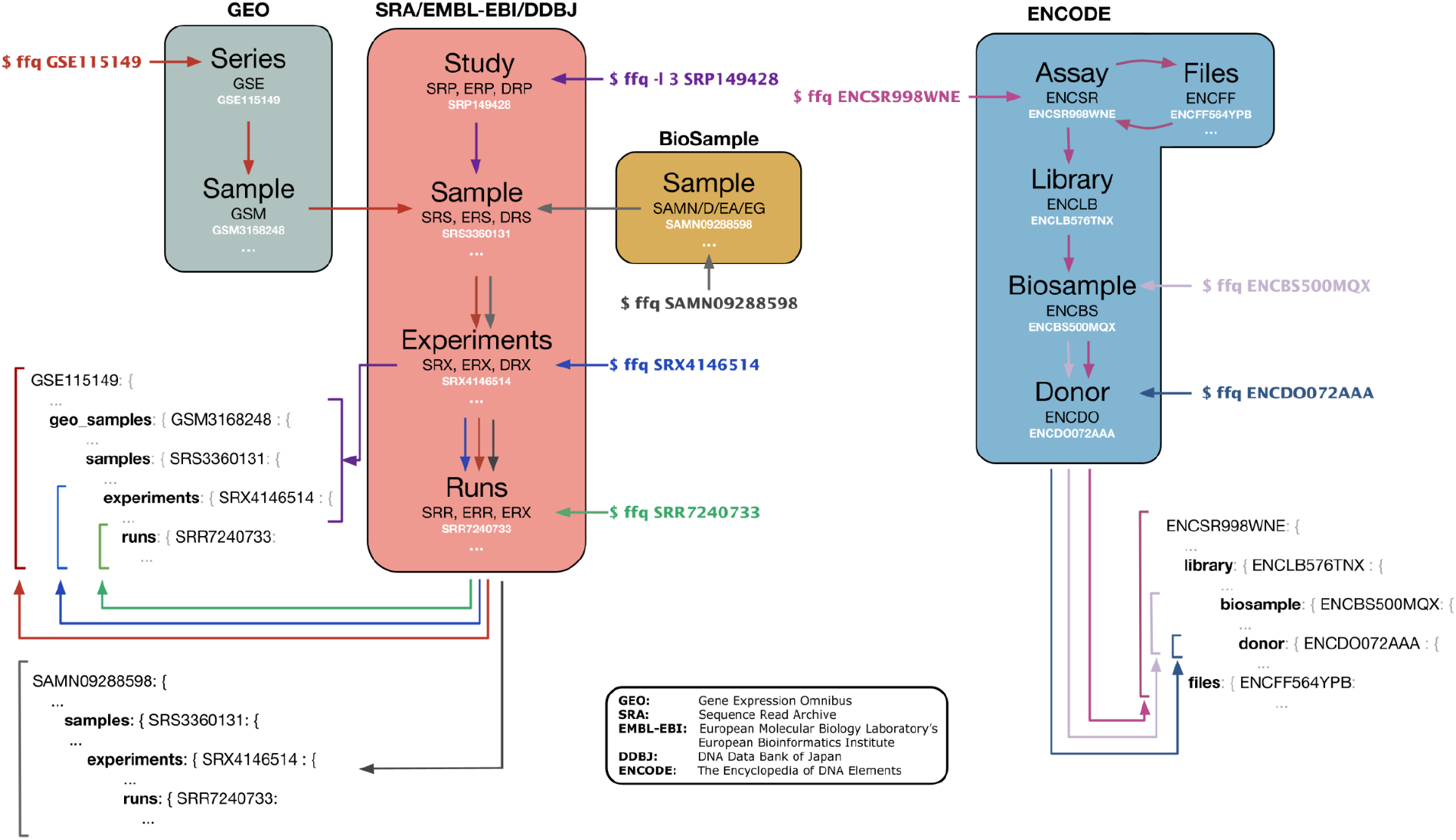
Metadata retrieval. *ffq* fetches and returns metadata as a JSON object by traversing the database hierarchy. Subsets of the database hierarchy can be returned by specifying -*l [level]*.

The SRA, ENA, and DDBJ databases all follow a similar hierarchical structure where studies are grouped into samples, experiments, and runs, a shared architecture that is useful and likely the result of the longstanding International Nucleotide Sequence Database Collaboration (INSDC) between the ENA, NCBI, and DDBJ. We note that the Genome Sequence Archive (GSA) (Chen *et al*., 2021; CNCB-NGDC Members and Partners, 2022) is not a member of the INSDC. However it also uses a similar hierarchical structure for its database, and regularly ingests data from the SRA, but does not expose its publicly available data for programmatic access.

The consistent database schemas used by members of the INSDC greatly simplifies metadata retrieval for *ffq*. For example, GEO accession codes are grouped hierarchically through Series and Samples and have external relations to SRA accession codes for raw sequencing data submitted to the SRA. This enables *ffq* to fetch metadata and processed data from GEO that submitters have associated with raw sequencing data stored in the SRA.

## Description

Based on the database architectures, we created *ffq* to fetch metadata using database accessions or paper DOIs as input. Importantly, *ffq* only fetches metadata and links to data files and does not offer data downloading. This deliberate design decision was motivated by the UNIX philosophy “Make each program do one thing well” (McIlroy *et al*., 1978).

The *ffq* options are summarized below:

- *ffq* [accession(s)]

○ Where [accession] can be any of the following: SR(R/X/S/P), ER(R/X/S/P), DR(R/X/S/P), GS(E/M), ENC(SR/BS/DO), CXR, SAM(N/D/EA/EG), DOI.
- *ffq* [-l level] [accession(s)]

○ Where [level] defines the hierarchy in the database to which data is subset data.
- *ffq* [--ftp] [--aws] [--gcp] [--ncbi] [accession(s)]

○ Where the flags correspond to the types of data-storage links for the raw data.
- *ffq* [-o out] [--split] [accession(s)]

○ Where [out] corresponds to a path on disk to save the JSON file and [--split] splits the metadata from multiple accessions into their own file.

Accession-based *ffq* metadata retrieval uses the NCBI’s Entrez programming utilities, ENA’s API, GEO’s FTP, and ENCODE’s API to programmatically access metadata with HTTP requests. DOI-based metadata retrieval first converts the DOI to the manuscript tile via the CrossRef API (Hendricks *et al*., 2020) and then retrieves all study accessions associated with the manuscript title with the ENA search API. Metadata is returned as a Javascript Object Notation (JSON) object. Run times for metadata retrieval vary depending on database up-time, server connection speed, and database rate-limiting, but generally we find that *ffq* can download metadata at a rate of 10s per sample. This rate includes short and deliberate delays we have added between HTTP requests to prevent a perceived Denial-of-Service.

## Usage and Documentation

The *ffq* tool is written in Python and can be installed with pip and conda. It has four dependencies and undergoes quality control via an automated testing framework that validates behavior against three Python versions (3.6, 3.7, and 3.8) covering 88% of the code. The JSON return objects make *ffq* interoperable with other tools such as *jq* for easy command-line parsing. Additionally, *ffq’s* modularity and simplicity make it extensible to other genomic databases. By leveraging existing APIs, *ffq* offers a lightweight solution for querying data that is guaranteed to be more up-to-date than tools that rely on regular database builds.

## Discussion

While *ffq* facilitates downloading of data from numerous genomic databases, the results retrieved are only useful to the extent that the metadata uploaded is meaningful and complete. Meaningful and complete user-generated data underlies the curation of genomic references essential for comparative genomic data analysis (Luebbert and Pachter, 2022). Unfortunately, there is little to no standardization of user-uploaded sequencing metadata (Wang *et al*., 2019; Rajesh *et al*., 2021), and metadata descriptions can become exceedingly complex for current multiplexed experiments where different assays with distinct data types are combined. Improvement of metadata uploading in machine-readable standard formats is essential if publicly available genomic data are to be usable by scientists in the future.

## Acknowledgments

This work was motivated by the need to obtain metadata for (Booeshaghi and Pachter, 2020). We thank Ali Mortazavi for his suggestion to include *ffq* querying of the ENCODE database.

